# BUSCO applications from quality assessments to gene prediction and phylogenomics

**DOI:** 10.1101/177485

**Authors:** Robert M. Waterhouse, Mathieu Seppey, Felipe A. Simão, Mosè Manni, Panagiotis Ioannidis, Guennadi Klioutchnikov, Evgenia V. Kriventseva, Evgeny M. Zdobnov

## Abstract

Genomics promises comprehensive surveying of genomes and metagenomes, but rapidly changing technologies and expanding data volumes make evaluation of completeness a challenging task. Technical sequencing quality metrics can be complemented by quantifying completeness in terms of the expected gene content of Benchmarking Universal Single-Copy Orthologs (BUSCO, http://busco.ezlab.org). Now in its third release, BUSCO utilities extend beyond quality control to applications in comparative genomics, gene predictor training, metagenomics, and phylogenomics.

---

Genomics approaches play a preeminent role in biological research because they are high-throughput and cost-effective, leading to the generation of ever-increasing volumes of data. This makes thorough quality control of sequencing data ‘products’, e.g. genomes, genes, or transcriptomes, ever more important. Addressing this, the Benchmarking Universal Single-Copy Ortholog (BUSCO) assessment tool provides intuitive quantitative measures of genomic data completeness in terms of expected gene content^1^. BUSCO identifies complete, duplicated, fragmented, and missing genes and enables like-for-like quality comparisons of different datasets. These features mean that BUSCO has become established as an essential genomics tool, using up-to-date data from many species and with broader utilities than the popular but now discontinued Core Eukaryotic Genes Mapping Approach^2^ (CEGMA). In this communication, we present the major BUSCO improvements, now in its third release as detailed below, with scenarios that highlight BUSCO’s wide-ranging genomics utilities: designed primarily for performing **genomics data quality control**, but also applicable for building robust **training sets for gene predictors**, selecting high-quality reference species for **comparative genomics analyses**, and identifying reliable markers for large-scale **phylogenomics and metagenomics studies**.

***Genomics data quality control*** motivated the delineation of the original BUSCO datasets^3^ and their subsequent integration with the assessment tool for analyzing the completeness of genome assemblies, annotated genes, and transcriptomes^1^. Benchmarking new genomes or gene sets against those of gold-standard model organisms or of closely-related species provides intuitive like-for-like comparisons. For transcriptomes, high completeness is expected for samples pooled from multiple life stages and tissues, while lower scores for targeted samples corroborate their specificity. Benchmarking can also help to guide iterative re-assemblies or re-annotations towards quantifiable improvements, e.g. the postman butterfly^4^ and Atlantic cod^5^. Here we assess three versions of the annotated chicken and honeybee genomes (see Methods), which have been the subject of extensive enhancements^6,7^ and clearly demonstrate the utility of BUSCO for quantifying successful improvements (Figure 1). Progressions from the initial, to intermediate, and latest versions of both species show improved completeness using the high-resolution hymenoptera or aves datasets and the lower-resolution metazoa dataset.

**Figure 1.**
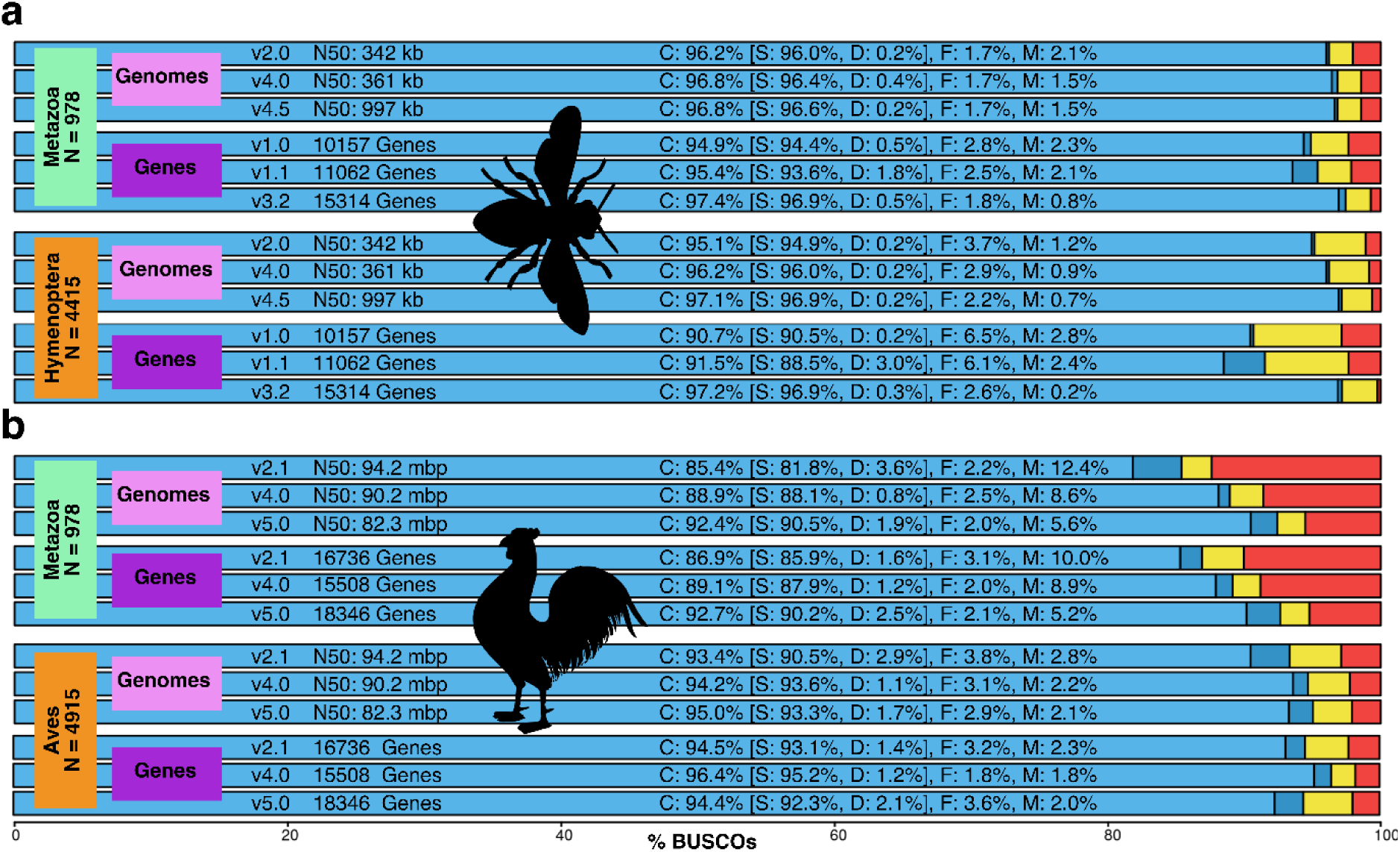
BUSCO completeness assessments for genomics data quality control. Assessments of initial, intermediate, and latest versions of the (**a**) honeybee and (**b**) chicken genomes and their annotated gene sets with the Metazoa, Hymenoptera, and Aves lineage datasets. Bar charts produced with the BUSCO plotting tool show proportions classified as complete (C, blues), complete single-copy (S, light blue), complete duplicated (D, dark blue), fragmented (F, yellow), and missing (M, red).

***Gene predictor training*** exemplifies BUSCO utilities beyond quality control, as gene models built during genome assessments represent ideal input data for parameterizations. Accurate prediction of protein-coding genes remains challenging, especially when supporting evidence such as homologs or native transcripts is not available and predictions are performed *ab initio*. This involves statistical modeling of nucleotide signatures and content to build gene models that best fit pre-trained parameter distributions. These vary considerably among species and thus require optimization, often employing high-quality gene annotations from native transcripts as input data. BUSCOs represent complementary predefined sets for such training procedures, without the need to perform RNA sequencing. Comparing Augustus^8^ predictions using BUSCO-trained parameters versus available pre-trained parameters from other species (see Methods) can show substantial improvements, e.g. BUSCO-trained *Strigamia* centipede, *Daphnia* waterflea and *Danaus* butterfly predictions are much better than using fruit fly parameters (Figure 2, Figure S1). Where species-specific-trained parameters are available, BUSCO training performs almost as well, e.g. tomato and thale cress, just as well, e.g. fruit fly and *Nasonia* wasp, or even better, e.g. *Tribolium* beetle (Figure 2, Figure S1). BUSCO employs Augustus for gene prediction so assessing genomes automatically generates Augustus-ready parameters trained on genes identified as complete. Additionally, the BUSCO-generated general feature format and GenBank-formatted gene models can be used as inputs for training other gene predictors like SNAP^9^. Running assembly assessments therefore provides users with high-quality gene model training data that can greatly improve genome annotation procedures.

**Figure 2.**
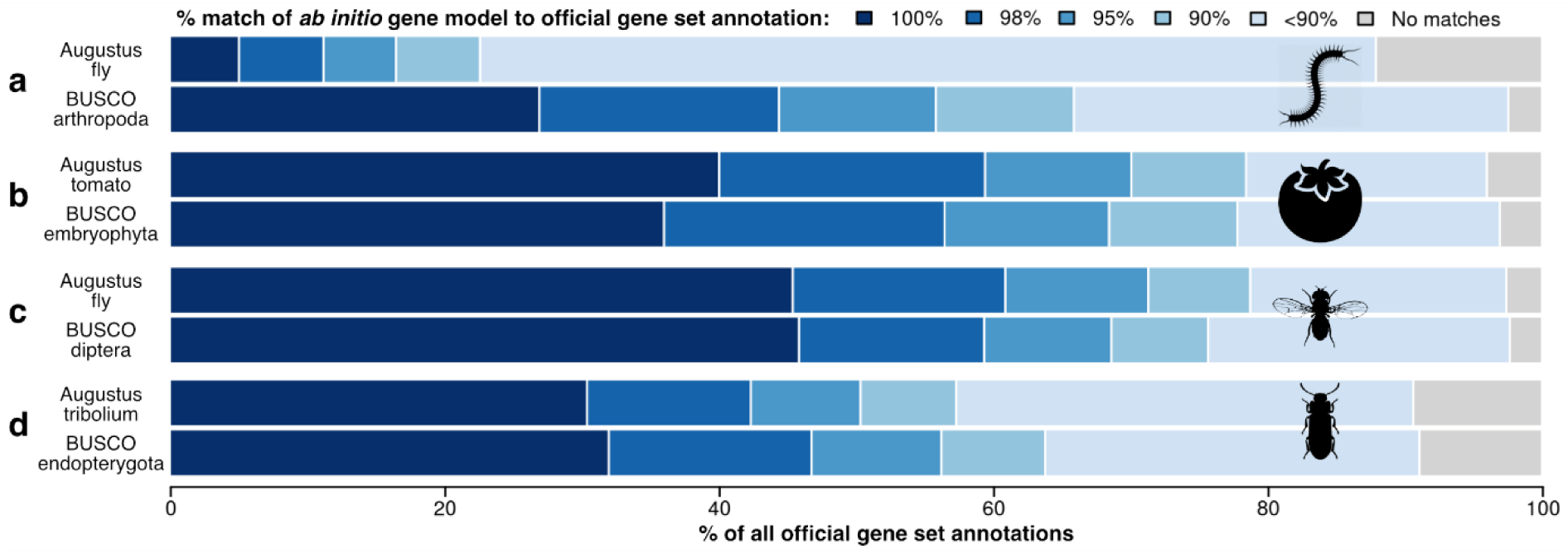
BUSCO-trained *ab initio* gene prediction with Augustus. When no pre-trained parameter set is available, e.g. for (**a**) the centipede, BUSCO-trained predictions are substantially better than using Augustus parameters from another arthropod (fly). Where species-specific-trained parameter sets are available, BUSCO-trained predictions are almost as good, e.g. (**b**) tomato, just as good, e.g. (**c**) fruit fly, or even better, e.g. (**d**) *Tribolium* beetle. Performance was assessed by computing the percent sequence length match of the *ab initio* gene models to the official gene set annotations for each species.

***Comparative genomics analyses*** are often sensitive to incomplete data, making the selection of high-quality datasets from representative species a critical first step for many studies. This becomes increasingly complex as the amount of available genomics data grows, especially as quality may vary considerably. Quantifying completeness can help to make objective selections, e.g. surveying 653 *Streptomyces* genomes identified the full complement of complete bacteria BUSCOs for only 63% of them^10^. Selecting those with the most genes does not guarantee quality, as genomes with many genes are not necessarily the most complete and those with fewer genes are not always less complete^11^. Selections will undoubtedly be influenced by considerations of taxonomic sampling, the availability of pertinent functional genomics data, the extent and/or accuracy of functional annotations, or simply historical usage. However, all else being equal, quantitative assessments with BUSCO offer logical selection criteria to help focus on the most complete genomic resources available. For example, assessing 135 *Lactobacillus* and 35 *Aspergillus* genomes and comparing these with their contiguity measures and total gene counts (see Methods) shows that RefSeq-designated references are not always the best available representatives (Figure S2). Comparing such metrics in this way therefore allows for the informed selection of the best quality representatives for subsequent comparative analyses.

***Phylogenomics*** takes advantage of whole genome or transcriptome data to reconstruct phylogenies that chart the relationships among organisms, a prerequisite for almost any evolutionary study. Being near-universal single-copy genes, BUSCOs represent predefined sets of reliable markers where assessments can identify shared subsets from different types of genomic data. For example, employing BUSCOs from insect genomes and transcriptomes to confirm Odonata-Neoptera relationships^12^, and from nearly 100 fungal genomes to reconstruct the Saccharomycotina phylogeny^13^. Analysis of seven rodent genomes and five transcriptomes illustrates the use of BUSCO to recover genes for phylogenetic inference (Figure 3). The identified genes were used to build a superalignment from which to estimate the species phylogeny (see Methods), which agrees with previous studies^14^. Assessments with the high-resolution Euarchontoglires or Mammalia datasets take longer but they identify more than three times as many universal single-copy markers than the lower-resolution Metazoa dataset. This illustrates the utility of BUSCO assessments to relatively quickly and easily identify reliable single-copy markers from different types of genomic data for phylogenomics analyses. Universal molecular markers are also essential in metagenomics studies, for phylogenetic classification of the surveyed microbiota, and where estimating relative abundances is greatly simplified if the markers are single-copy^15^. Hence BUSCOs also represent ideal markers for applications in metagenomics.

**Figure 3.**
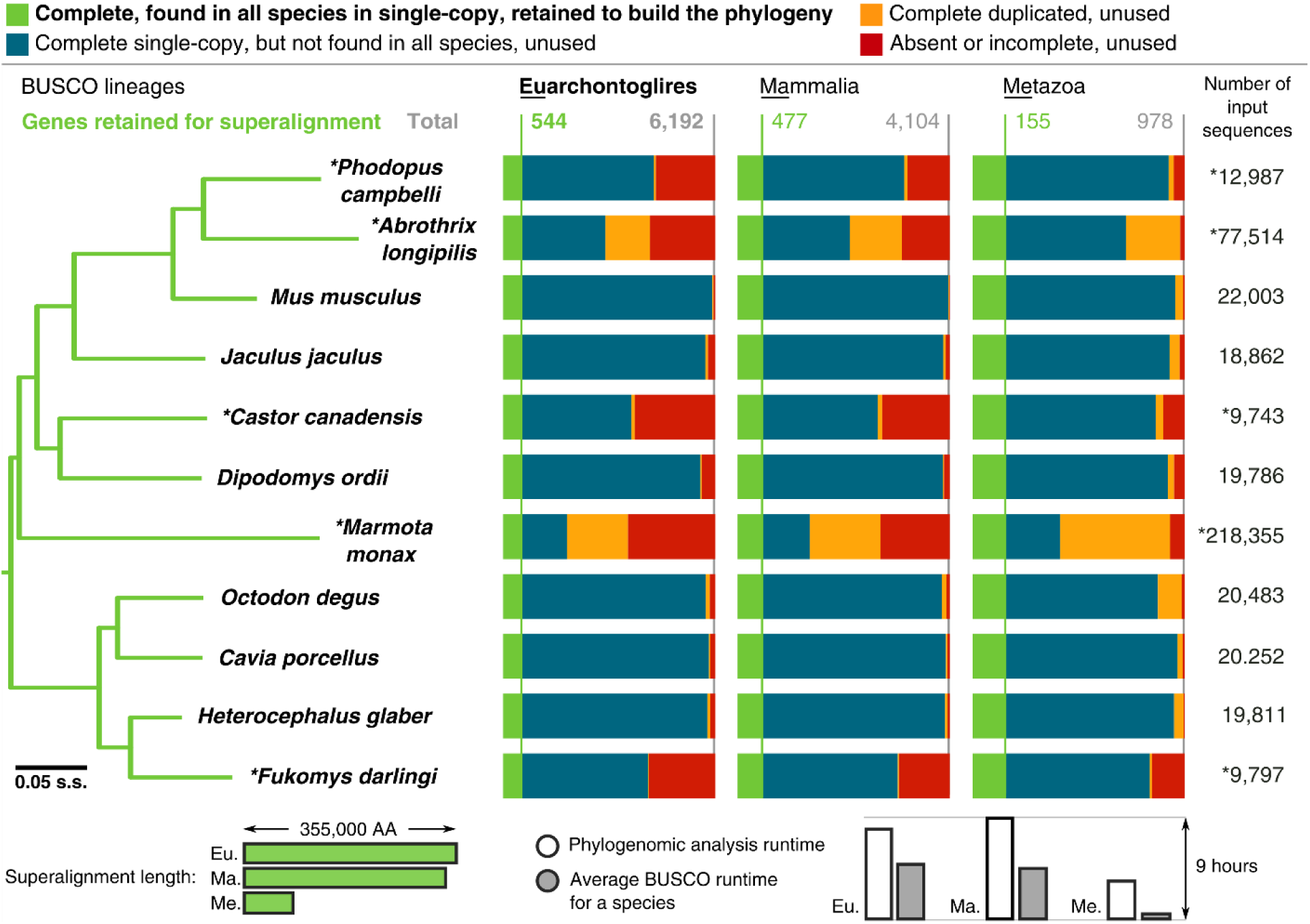
Genome and transcriptome BUSCO assessments to identify universal single-copy markers for phylogenomics studies. The phylogeny was generated using the Euarchontoglires results to identify complete single-copy orthologs found in all species for building the superalignment used for maximum likelihood tree reconstruction (see Methods). Mammalia and Metazoa results produced identical tree topologies. Bars below the BUSCO results show how the sizes of the assessment datasets influence the superalignment lengths and the analysis runtimes. The tree was rooted with the rabbit, all nodes have 100% bootstrap support, branch lengths are in substitutions per site (s.s.).

BUSCO datasets comprise genes evolving under ‘single-copy control’^16^, i.e. within each lineage they are near-universally present as single-copy orthologs. This property underlies the evolutionary expectation that they should be present, and present only once, in a complete assembly or gene set. Completeness is quantified in terms of this expected gene content by assessing the orthology status of predicted genes using BUSCO sequence profiles. BUSCOs are carefully selected with finely-tuned score and length cut-offs that maximize precision and recall, but as both gene prediction and orthology assignment are challenging tasks, assessments may still fall short of 100% correct classification. Additionally, while input species selection explicitly avoids over-sampling closely-related species, the choices must be made from currently available resources that are not phylogenetically evenly distributed. With these caveats in mind, BUSCO offers like-for-like assessments for genomics data quality control, which perform well in qualitative comparisons with alternative measures. For example, metrics based on genome alignments that quantified completeness of ultraconserved elements and protein-coding exons by comparing 20 vertebrates to human^17^ showed overall very good agreement with BUSCO results. Furthermore, assessing 12 plants^18^ with BUSCO, CEGMA, core plant Gene Families, and Expressed Sequence Tag mapping also showed good agreement. BUSCO therefore offers reliable measures of completeness that agree with alternative approaches, are applicable to different genomic data types, and offer like-for-like comparisons. This utility extends to additional genomics applications including defining datasets for training gene predictors, facilitating objective selection of representatives for comparative studies, and identifying reliable markers for phylogenomics and metagenomics.

Since the initial BUSCO release, development has aimed to address user needs with BUSCO v2 implementing improvements to the underlying analysis software as well as updated and extended datasets covering additional lineages based on orthologs from OrthoDB v9^19^. To facilitate high-throughput assessments, BUSCO v3 implements a refactoring of the code to make it more flexible and extendable by simplifying installation and introducing control through a configuration file. Additionally, visualization of the results is enabled with a plotting tool that generates easily configurable bar charts. The software is distributed through GitLab, it is also available as an Ubuntu virtual machine, and it has been integrated as an online service for logged-in users at http://www.orthodb.org. These and other new features, options, setup instructions, as well as best practices are all described in detail in the updated user guide (http://busco.ezlab.org). With many more new species being sequenced, future BUSCO releases will focus on adding new lineages for which species sampling becomes rich enough to build reliable datasets as well as providing higher resolution with larger lineage-specific datasets.

## METHODS

Methods, including statements of data availability and any associated accession codes and references, are available in the supplementary online material.

## ACKNOWLEDGEMENTS

The authors would like to thank all members of the Zdobnov laboratory and our enthusiastic users who have made suggestions to improve the codebase, requested new lineage-specific datasets, and beta-tested the BUSCO updates. This work was partly supported by the Swiss Institute of Bioinformatics SER funding, University of Geneva funding, and the Swiss National Science Foundation grant 31003A_143936 to E.M.Z., R.M.W. was supported by Swiss National Science Foundation grant PP00P3_170664. Some of the computations were performed at the Vital-IT (http://www.vital-it.ch) center for high-performance computing of the Swiss Institute of Bioinformatics.

### AUTHOR CONTRIBUTIONS

E.M.Z., E.V.K. and R.M.W. conceived the study. F.A.S., G.K., M.S., and R.M.W. developed the software, collated datasets, and performed the analyses. M.M. and P.I. contributed to dataset collation and software testing. E.M.Z., E.V.K., F.A.S., M.S., and R.M.W. wrote the manuscript with input from all authors.

### COMPETING FINANCIAL INTERESTS

The authors declare no competing financial interests.

